# EPHierStats: a statistical tool to model the hierarchical relationships in electrophysiological data

**DOI:** 10.1101/2022.03.23.485501

**Authors:** Jenya Kolpakova, Jonathan D. Marvel-Zuccola, Kensuke Futai, Gilles E. Martin, Vincent van der Vinne

**Affiliations:** Brudnick Neuropsychiatric Research Institute, Department of Neurobiology, University of Massachusetts Chan Medical School, Worcester MA 01655, USA, Worcester MA 01655, USA; Graduate Program in Neuroscience, Morningside Graduate School of Biomedical Sciences, UMass Chan Medical School, Worcester MA 01655, USA, Worcester MA 01655, USA; Biology Department, Williams College, Williamstown MA 01247, USA

## Abstract

Electrophysiological datasets are typically analyzed under the assumption that repeated measurements of the same unit of analysis (i.e. neuron or animal) can be treated as statistically independent. Recently, this assumption has been questioned and our data confirms and quantifies this skepticism using *ex vivo* slice recordings of synaptic currents in D1R^+^ medium spiny neurons in the nucleus accumbens. We therefore present EPHierStats as a statistical framework to analyze electrophysiological datasets with large numbers of measurements (>100) per unit of analysis. This novel analysis framework enables encoding of the full hierarchical relationships between measurements in a mixed-effects general linear model while also analyzing the distribution of values in assessed variables. Our method can easily be adapted to analyze a wide range of repeated-measures electrophysiological experiments. Implementation of the EPHierStats tool will aid the adaption of modern statistical approaches that prevent pseudoreplication and its associated false discovery rate while enabling statistical assessments of the complex relationships inherent to the field of neuroscience.

## Introduction

Electrophysiological recordings of the activity of neurons provide insight into the functioning of the brain at the level of its fundamental components (Kandel et al., 2021). Electrophysiological recordings of individual cells can be performed using a suite of different approaches, including the recording of action potentials, single-channels, excitatory and inhibitory postsynaptic currents (EPSCs/IPSCs), and/or multi-electrode array recordings. All of these approaches can record the electrophysiological behavior of individual neurons and do so by performing (many) repeated measurements of each neuron. The repeated nature of electrophysiological measurements does however complicate the analysis of these recordings and commonly used analysis approaches are often overly conservative or unreliable due to pseudoreplication [Aarts et al., 2014; Yu et al., 2022].

One of the most common analyses for electrophysiological datasets is to assess whether the average of an output parameter is different between treatment conditions. Since repeated measurements from the same neuron will typically be more alike compared to measurements from a different neuron (thus violating the homogeneity of variance assumption), analyses often calculate the average output parameter per cell and statistically analyze these average values [Aarts et al., 2014]. Although this approach circumvents the problem of pseudoreplication, the resulting analysis will suffer from an unnecessarily low statistical power. Also, with regression to the mean inherently occurring in cells with more measurements, cells that were sampled less often will be more likely to have an extreme value [Stigler, 1997]. Due to the normalizing at the level of individual cells, an outsized weight will thus be attached to these more extreme values. Finally, it is important to recognize that the distribution of electrophysiological values typically resembles a Poisson distribution, thus requiring either a non-parametric statistical test or transformation of the data to ensure that the residuals are normally distributed [Limpert et al., 2001; Yu et al., 2022].

Another research question often addressed using electrophysiological measurements is whether the distribution of values is altered between treatments. An example of such an altered distribution can be observed in a dataset where one treatment results in an increase of activity at a specific part of the distribution of outcomes [Mao et al., 2018]. A typically employed statistical test to assess changes in the distribution of values is the two-sample Kolmogorov-Smirnov test [Massey, 1951; Manabe et al., 1992]. This test does however not account for the specific neuron that each measurement was taken from, and the test thus requires that the distribution of values is the same in each of the neurons included in a group. Conversely, in cases where the distribution of measurement values is dependent on the neuron from which the recording was taken (or another similarly confounding factor), the repeated measurements within those neurons cannot be considered as statistically independent and thus pseudoreplication will result in an overestimation of the statistical effect size. Finally, the Kolmogorov-Smirnov test only assesses the value at which the two samples are most divergent and is thus incapable of assessing statistical differences at any other part of the distribution.

This paper first quantifies the statistical influences of the neuron, slice, animal, and experimenter levels on individual EPSC measurements of D1-receptor^+^ medium spiny neurons in the nucleus accumbens of mice. These analyses of EPSC frequency and amplitude illustrate the necessity of incorporating the underlying hierarchical relationships in the statistical analysis of such an electrophysiological dataset. Subdivision of the repeated measurements of each neuron into multiple subsequent intervals provides a way to generate biological replicates that increase statistical power while also enabling assessment of the distribution of outcomes. We present a flow scheme to implement this analytical approach to a hierarchical electrophysiological dataset and demonstrate its utility by analyzing an existing dataset. Overall, this novel statistical framework enables the use of powerful mixed-effects general linear modeling approaches to analyze the complex relationships that are common in electrophysiological datasets.

## Materials & Methods

### Electrophysiological measurements

All animal experiments used 8-10 weeks old male C57Bl/6J Drd1a-tdTomato mice and were approved by the Institutional Animal Care and Use Committee of the University of Massachusetts Medical School. Mice were maintained at constant temperature (22 ± 1 °C) and humidity in a 12 h:12 h light–dark cycle with water and food available *ad libitum*. Mouse acute brain slices were prepared according to methods previously described (Kolpakova et al, 2021). Whole-cell patch clamp recordings of spontaneous EPSCs were acquired from dopamine-1 receptor^+^ (D1R) MSNs in the NAc. EPSCs were isolated by recording in the presence of 15 μM Bicuculine (GABA receptor antagonist). Spontaneous EPSCs were acquired for 4-6 mins using gap-free recording at MSN resting membrane potential (Kolpakova et al, 2021). EPSCs’ amplitude and interevent interval were determined using Clampfit (pClamp 11 software suite, Molecular Devices).

### Data Analysis and Statistics

The interevent interval and amplitude of all EPSCs identified by Clampfit were associated with the corresponding mouse, experimenter, brain slice, neuron, and time interval for each individual measurement. For each neuron, EPSCs occurring during three minutes of recording were included and subdivided into three one-minute time intervals for statistical analyses. Based on visual assessments of the distribution of residuals resulting from different transformations (^n^✓x, with n = [1:400]) we concluded that the ^40^✓x transformation was adequate and enabled the use of parametric statistical approaches. The partitioning of variance was assessed separately for interevent interval and amplitude in a series of mixed-effects general linear models incorporating all the independent variables listed above as random variables to determine the variance explained. The statistical significance of individual variables was assessed in a series of models with all variables included as random factors except for the assessed variable. In all these analyses, variables were nested to reflect the hierarchical relationships between biological levels of organization. All statistical analyses were performed using SAS JMP 7.0.

Assessments of the distribution of values were performed using different approaches to describe the distribution. In the traditional approach, all untransformed values were first averaged per neuron to describe the average response, while a cumulative distribution was calculated while ignoring the neuron from which each measurement was taken. Our novel approach first visualized the distribution of values in a subset of seven neurons taken from separate mice and measured by the same experimenter. The value closest to the 5^th^, 25^th^, median, 75^th^ and 95^th^ percentile was determined separately for each of the three-minute sets of measurements in each neuron with means ± SEM calculated based on transformed values. The subsequent visualization of percentile values in individual neurons determined the value closest to the 5^th^, 25^th^, median, 75^th^ and 95^th^ percentile separately in each of the three one-minute intervals available in each neuron and presents the mean ± SEM based on the transformed values of these three values.

### *EPHierStats* procedure

The EPHierStats approach aims to optimize a tradeoff between the most powerful statistical analysis of average responses and the ability to describe the distribution of outcomes. Subdividing measurements into biological replicates of 25 observations enables the selection of values corresponding exactly to the 6^th^, 26^th^, median, 74^th^, and 94^th^ percentile values. These values were selected to represent the median, shoulders, and extreme values of the distribution we aimed to describe but could easily be replaced with other percentile values if appropriate. The inclusion of ten replicates of 25 measurements per condition enabled very powerful statistical comparisons but a lower number of replicate intervals will typically be sufficient to accurately quantify within-neuron variability. Selection of the five values per interval was accomplished here in MS Excel following organization of the dataset using the Sort command and can be accomplished easily in a wide range of software packages. Visual inspection of residuals of the various models confirmed that normalization using the ^40^✓x transformation worked well for our dataset and enabled us to use parametric statistical tests. Proper encoding of the hierarchical relationships between the measurements from biological replicates is done using a mixed-effects general linear model in which (at a minimum) neuron and time interval should be included as random variables. The percentile [6, 26, 50, 74, 94] of each included value should always be included as a categorical fixed main effect in the statistical model as well as the interaction between the percentile and the factor of interest (i.e. percentile*genotype). If this interaction term does not substantially reduce the unexplained variance in the model, the interpretation of the statistical results might be simplified by removing this interaction term from the model.

#### Validation of the EPHierStats analysis approach

The utility of the EPHierStats approach was assessed by re-analyzing a previously published dataset [Mao et al, 2018] that tested the involvement of the Chromatin reader L3mbtl1 in synaptic scaling induced by pharmacological treatments [Turrigiano, 2011]. In line with traditional approaches to analyze a dataset like this, our initial statistical comparisons analyzed the average miniature EPSC (mEPSC) amplitude for each neuron. A second model described the hierarchical relationships between different neurons more extensively by adding Mouse ID as a random variable to the initial model. The final statistical model applied the EPHierStats approach outlined above and included Mouse ID, Neuron ID, and Time interval as random variables. All statistical comparisons were made using mixed-effects general linear models with genotype, pharmacological treatment, and their interaction included in all models as fixed effects. The EPHierStats model also included percentile and its interactions as fixed effects. The criterion for statistical significance was *p*<0.05 for all experiments and post-hoc comparisons were made using Tukey-HSD tests.

## Results

### Repeated electrophysiological measurements are not statistically independent

Common analysis methods assessing changes in the average electrophysiological response of neurons to experimental treatments often assume that the repeated measurements taken from multiple neurons are statistically independent. Statistical independence is a core assumption underlying all statistical tests and it requires that the residual of one or a subset of datapoints cannot be used to predict the residual of any other datapoints [Grafen & Hails, 2002]. The assumption of statistical independence is typically violated when repeated measurements are performed since measurements from the same neuron are likely to be more similar compared to measurements from a different neuron. To restore statistical independence in the analysis of a repeated-measurements dataset, it is necessary to properly encode the hierarchical relationships between measurements in the statistical analysis (e.g. repeated-measures ANOVA). Beyond the expected similarities between individual neurons, measurements taken from neurons in the same brain slice and/or the same animal might similarly result in violations of the assumption of statistical independence.

To establish which levels of biological organization should be incorporated in the statistical analysis of repeated electrophysical measurements in order to ensure statistical independence, repeated EPSC measurements were performed by 2 experimenters from 110 neurons on 36 brain slices derived from 8 mice (Fig. 1). As expected, both the inter-event interval and amplitude of these EPSCs were Poisson distributed (Fig. 1A, C), thus requiring transformation (^40^✓x) to enable the use of parametric statistical approaches (Fig. 1B, D). Subdividing the recordings into one-minute intervals provided three biological replicates of repeated EPSCs for each neuron that illustrated the low within-neuron variability relative to the between-neuron variability of the EPSCs’ mean interevent interval and amplitude (Fig. 1E, F). Partitioning of the variance in interevent interval between the different levels of biological organization showed that the neuron ID and slice ID could explain 18.6 % (F_74,196.3_ = 34.44, p < 0.0001) and 9.5 % (F_28,38.02_ = 3.916, p < 0.0001) of the total variance, respectively (Fig. 1G). The time interval (0.9 %, F_219,49592_ = 2.856, p < 0.0001), experimenter (1.5 %, F_1,32.33_ = 2.562, p = 0.1192), and mouse (1.9 %, F_7,25.1_ = 1.413, p = 0.2438) only marginally influenced the interevent interval duration. Analyzing the distribution of variance in EPSCs’ amplitude demonstrated that neuron ID explained 23.8 % (F_74,225.1_ = 49.52, p < 0.0001) of variance, while slice (−0.9 %, F_28,1_ = 2.092, p = 0.5050), interval (0.7 %, F_219,50002_ = 2.423, p < 0.0001), experimenter (2.6 %, F_1,16.66_ = 9.078, p = 0.0080), and mouse (−0.7 %, F_7,18.06_ = 0.4903, p = 0.8294) barely affected amplitude (Fig. 1I). In summary, these results show that repeated EPSC measurements are not statistically independent and that the neuron from which recordings are taken should be taken into account when analyzing this kind of electrophysiological recordings.

**Figure 1:**
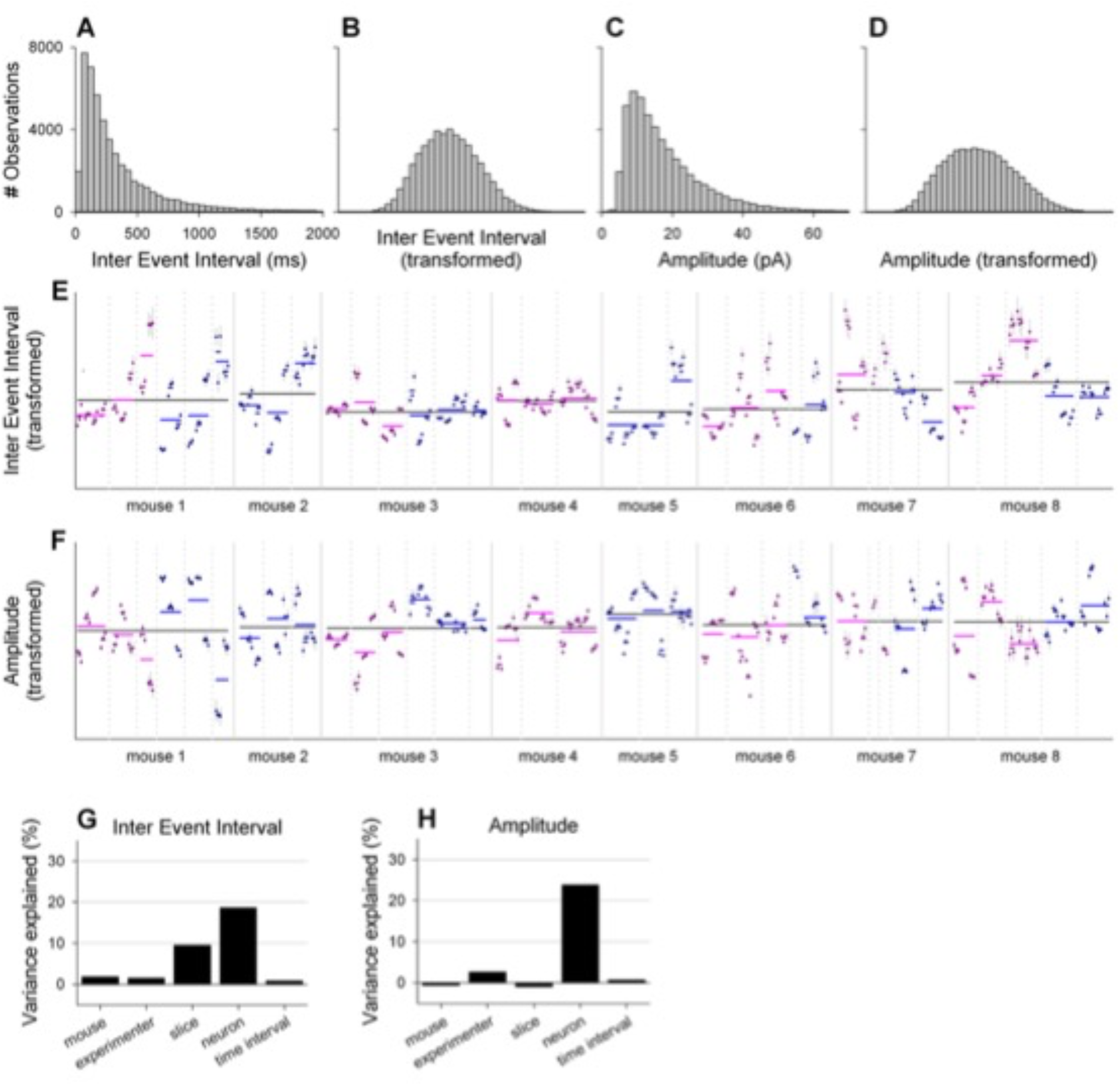
Hierarchical relationships shape measurements of EPSCs interevent interval and amplitude. (A, B) Histograms of raw (A) and transformed (^40^✓x, B) EPSCs interevent interval measurements. (C, D) Histograms of raw (C) and transformed (^40^✓x, D) EPSCs amplitude measurements. (E) Average ± SEM of transformed EPSCs interevent intervals for three one-minute intervals per neuron. Measurements taken from neurons on the same brain slice are separated by dotted vertical lines while continuous vertical lines separate mice. Recordings taken by two different experimenters are separated by color (red and blue). Averages per neuron (dark colored), slice (light colored), and mouse (black) are indicated by horizontal lines. (F) Average ± SEM of transformed EPSCs amplitudes for three one-minute intervals per neuron. Drawing conventions are the same as in (E). (G, H) Proportion of variance in ESPCs interevent interval (G) and amplitude (H) explained by different model components.

### Analyzing distributions of electrophysiological measurements

Describing the factors that influence EPSCs is a key element of most electrophysiological studies. Such characterizations are typically accomplished through a two-step process that describes both the overall average as well as the distribution of values of the parameter being measured. Comparisons of the average response of a group of neurons is traditionally done by describing the average response of each neuron using a single parameter, typically the mean or median response, and statistically analyzing these values (Fig. 2A, G). Beyond assessing the average changes in interevent interval and amplitude, electrophysiological studies often assess the whole distribution of values to identify whether experimental manipulations specifically affect a subset of frequencies and/or strengths of EPSCs. Traditionally, such an analysis is performed by visually assessing the cumulative frequency probability curves of all measurements in a specific condition (Fig. 2B, H). Statistically, this visual approach is supported by a two-sample Kolmogorov-Smirnov test that assesses whether the maximal difference between two curves is larger than a statistical threshold [Massey, 1951]. This statistical comparison does however ignore all points of sub-maximal divergence and does not account for between-neuron differences.

**Figure 2:**
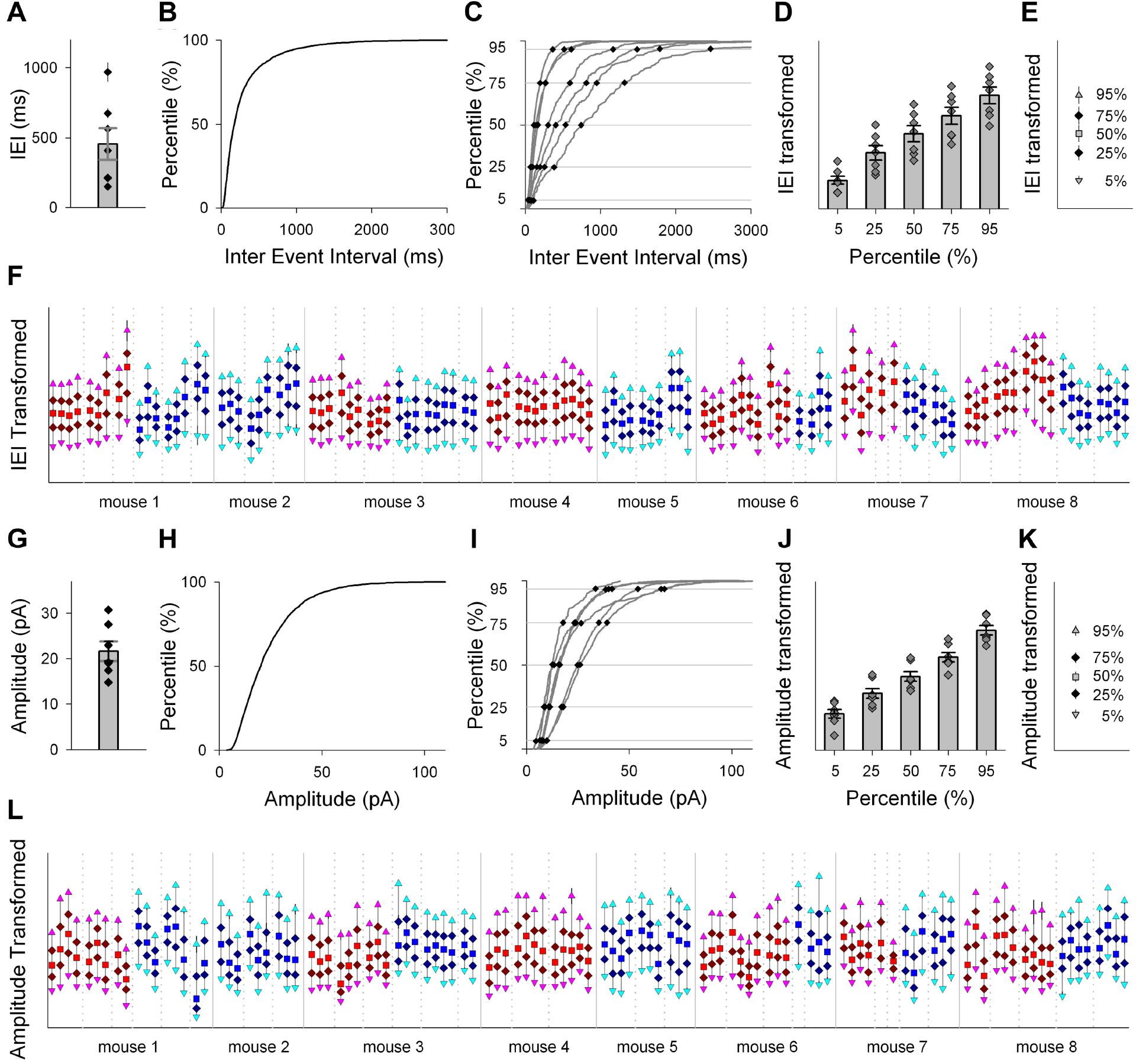
Describing the distribution of EPSC measurements. (A, G) Bar graph representing the traditional method to depict average responses showing group mean ± SEM of ESPCs interevent interval (A) and amplitude (G) with individual neurons (mean ± SEM) plotted as dots. Data obtained from seven individual neurons is included (left most neuron of each mouse analyzed by experimenter Blue in Fig. 1E; same in elements B/H, C/I, D/J, and E/K). (B, H) Line graph representing the traditional method to depict the distribution of measurement values in a cumulative frequency distribution plot of the EPSCs interevent interval (B) and amplitude (H). (C, I) Cumulative frequency distribution curves of EPSCs interevent interval (C) and amplitude (I) calculated separately for each of the seven included neurons (gray lines). Black dots represent the EPSCs interevent interval (C) and amplitude (I) of individual neurons at the indicated percentiles. (D, J) Bar graphs (mean ± SEM) of transformed (^40^✓x) EPSCs interevent interval (C) and amplitude (J) for five investigated percentiles. Dark gray dots (mean ± SEM) depict values for each of the seven neurons and correspond to black dots in C/I. (E, K) Mean ± SEM of transformed EPSCs interevent interval (E) and amplitude (K) at the indicated percentiles for seven neurons. The five points correspond to the bars in D/J. (F, L) Distribution of transformed EPSCs interevent interval (F) and amplitude (L) measurements of all neurons as described by five selected percentiles. Points represent the mean ± SEM of the specified percentile value determined separately for the first-, second-, and third-minute recording of each neuron. Measurements taken from neurons on the same brain slice are separated by dotted vertical lines while continuous vertical lines separate animals. Recordings taken by two different experimenters are separated by color (red and blue).

Describing the distribution of measurements separately for each neuron enables both the partitioning of variance between the hierarchical levels shaping measurement outcomes as well as the statistical assessment of the whole distribution of values. In practice, such a comparison becomes very complicated to interpret when each of a large number of observations (>100) is compared between neurons. We propose to calculate the 5^th^, 25^th^, 50^th^, 75^th^, and 95^th^ percentile values for each neuron as a way to describe the distribution of measurements for each neuron in sufficient detail to be both meaningful and comprehensible (Fig. 2C, I). These values represent the median, shoulders, and extreme values of the distribution of responses. When transformed (^40^✓x), the residuals at all the five assessed percentiles have comparable values and are normally distributed (Fig. 2D, J), thus enabling statistical comparisons using parametric statistics. Plotting the group averages of each of the percentiles provides a convenient way to visually represent the average distribution of measurement outcomes (Fig. 2E, K). Applying this visualization approach further illustrates the substantial variability between cells in both interevent interval (Fig. 2F) and amplitude of EPSCs (Fig. 2L), in line with the analysis of the same data presented in Figure 1E and 1F.

All statistical analyses are based on the partitioning of variance between underlying independent variables. Statistical power can typically be increased by including biological replicates and the repeated measurements that are common in electrophysiological studies provide a convenient way to do so. As outlined above, the distribution of measurement outcomes can be described by the 5^th^, 25^th^, 50^th^, 75^th^ and 95^th^ percentile values. Doing so based on the ∼250 measurements that our lab typically uses to describe a neuron’s characteristics provides a highly precise value describing the distribution of each individual neuron. However, from a statistical perspective, it is more powerful to subdivide this relatively large set of measurements into multiple biological replicates that, although each individually less precise, enables a more thorough partitioning of the variance in our dataset. We feel that subdivision into intervals of 25 events provides a sweet spot where the distribution of values can be meaningfully described with five percentile levels while the ten intervals allow for the estimation of within-neuron variance in these five values (Fig. 3). This then allows separation of the substantial (see Fig. 1) within-neuron variance from the unexplained variance, thus making statistical comparisons between neurons and especially within-neuron comparisons of experimental conditions much more powerful.

**Figure 3:**
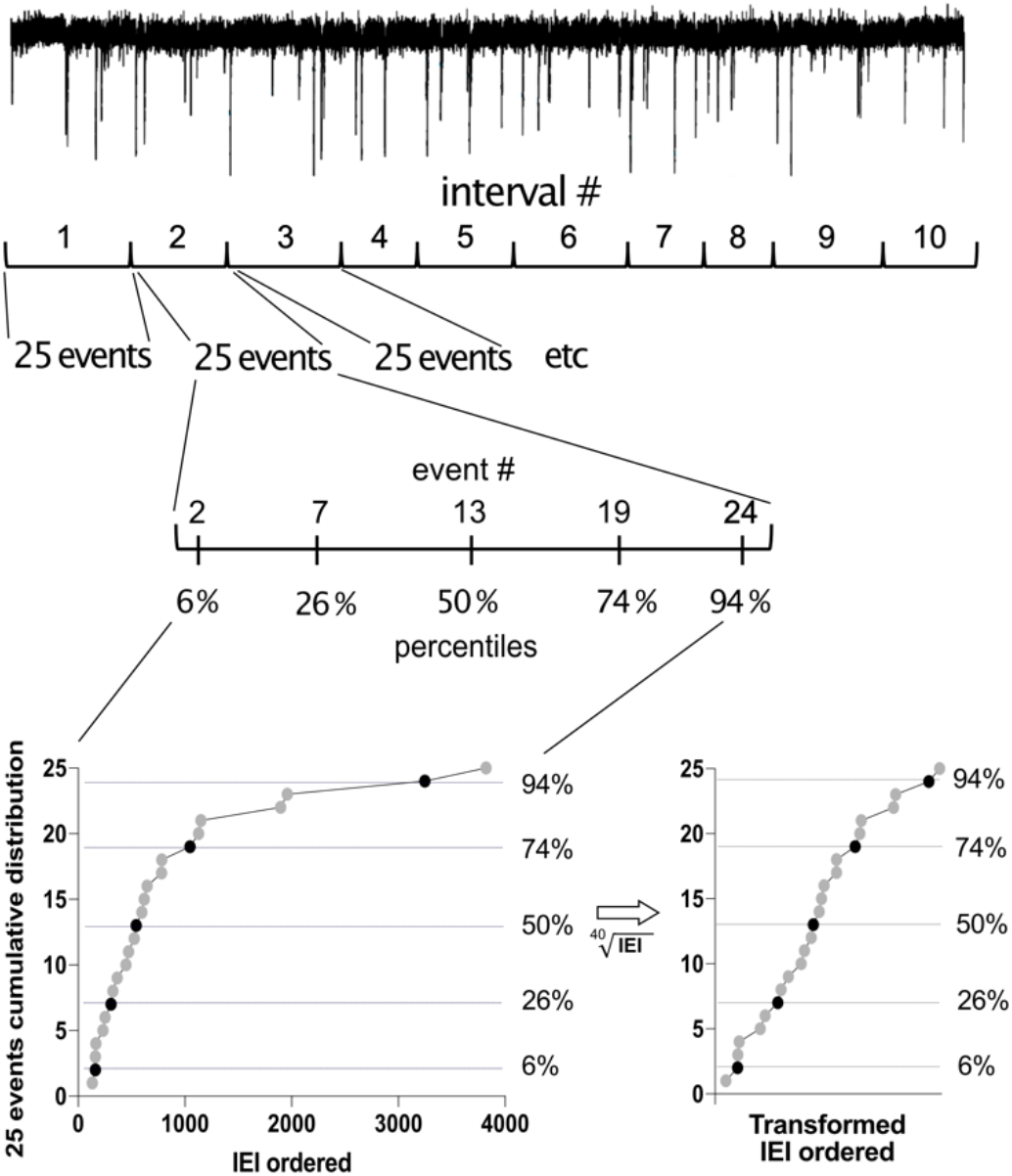
Partitioning of EPSCs raw recording events for statistical analysis. Recording of 250 EPSC inter-event intervals is divided into 10 biological replicates of 25 events. Each replicate set of 25 values is ordered from smallest to largest and the 2^nd^ (6^th^ percentile), 7^th^ (26^th^ percentile), 13^th^ (50^th^ percentile), 19^th^ (74^th^ percentile), and 24^th^ (94^th^ percentile) are taken. Together these values describe the distribution of measurement values, representing the median (50%), shoulders (26 and 74%) and extremes (6 and 94%) of the distribution. Transformation of the Poisson-distributed raw values typically enables statistical analyses using parametric tests such as mixed-effects general linear models.

### EPHierStats enables exposure of complex relationships in electrophysiological data

The utility of our novel statistical approach describing repeated electrophysiological measurements as a distribution of outcomes in a hierarchical statistical model was tested by re-analyzing a previously published dataset [Mao et al., 2018]. This dataset tested the involvement of the Chromatin reader L3mbtl1 in synaptic scaling that compensate for activity perturbation and maintain the excitatory and inhibitory balance. Hippocampal primary neurons were prepared from wildtype and L3mbtl1 KO mice and synaptic scaling was induced by applying picrotoxin (PTX) or tetrodotoxin (TTX) that increased or decreased neuronal activity, respectively. Miniature EPSCs were recorded 48 hours after induction of synaptic scaling.

The original analysis of this data was done by performing a series of two-sample Kolmogorov-Smirnov tests comparing pharmacological treatments with controls within both genotypes (Fig. 4A). This showed that in wildtype mice, PTX and TTX treatment both altered at least some part of the distribution of mEPSC amplitudes, while in L3mbtl1-KO mice only TTX pre-exposure significantly shifted some part of the distribution of mEPSC amplitudes. Unfortunately, these statistical tests provide only very limited information about what parts of the distribution are affected, since the Kolmogorov-Smirnov test only assesses the amplitude with maximal divergence (indicated in Fig. 4A by the vertical lines connecting the compared curves) and does not assess any other part of the distribution. Furthermore, the Kolmogorov-Smirnov test does not account for the clustering of repeated measurements from the same neuron (see Fig. 1F, I) and instead treats all measurements as statistically independent, thus likely overestimating the effect size and increasing the chance of making a type I error.

**Figure 4:**
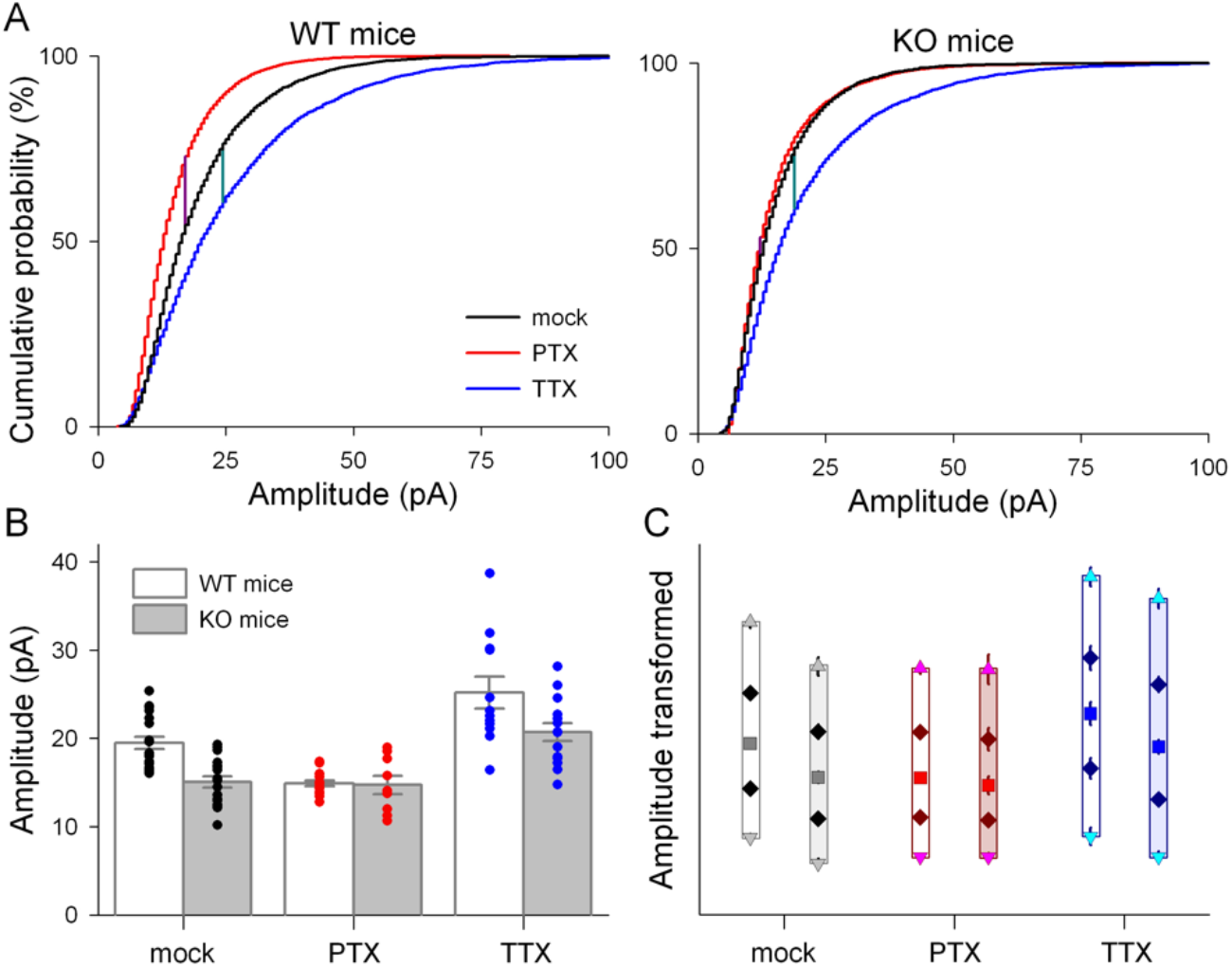
Application of EPHierStats exposes relationships in electrophysiological dataset. (A) Cumulative frequency distribution plots of mEPSC amplitude for wildtype (left) and L3mbtl1-KO mice (right) representing the traditional method for assessing differences in the distribution of measurements. Cumulative frequency distribution plots are depicted for all measurements from all neurons in each of the three treatment groups (mock, PTX, TTX). Vertical lines connecting cumulative distribution curves represent the test amplitude and the maximal percentile difference assessed by the two-sample Kolmogorov-Smirnov tests. (B) Bar graphs representing the mean ± SEM mEPSCs amplitude for six groups defined by genotype and pre-exposure treatment. Dots represent the mean mEPSCs amplitude for individual neurons. (C) The effects of L3mbtl1 genotype and pharmacological pre-treatment are most pronounced on mEPSCs with a larger amplitude. The 6^th^, 26^th^, 50^th^, 74^th^, and 94^th^ percentile of mEPSCs amplitudes is depicted for each genotype and pre-treatment combination (mean ± SEM). Colors represent pharmacological pre-treatment and bar opacity indicates genotype (white: wildtype, darker: L3mbtl1-KO).

The effects of incorporating the hierarchical nature of electrophysiological measurements into the statistical assessment of the average change in mEPSC amplitude (Fig. 4B) were analyzed by comparing three statistical models that became progressively more complex below. In line with the most common approach to analyze this type of dataset, model one described the response of each neuron by the mean amplitude and assumed that all neurons were statistically independent units. Based on this approach, the conclusion that the pharmacological treatment of neurons has divergent effects depending on the genotype of the mice would be justified (Genotype*Pharmacological treatment: F_2,81_ = 3.121, p = 0.0495). However, the second model showed that accounting for the relationships between neurons by adding the animal from which each neuron was derived as a random variable to the statistical model reduces both the degrees of freedom and the effect size. As a result, this model did not support the conclusion that the pharmacological treatment of neurons has divergent effects depending on the genotype of the mice (Genotype*Pharmacological treatment: F_2,73.4_ = 2.657, p = 0.0769).

The third statistical assessment of how the presence or absence of L3mbtl1 altered the mEPSC amplitude in response to PTX and TTX leveraged the repeated measures taken from each neuron and used the EPHierStats approach outlined in Figure 3 to generate biological replicates describing the distribution of mEPSC amplitudes. The hierarchical relationships between these repeated measurements were incorporated into the model by including the mouse, neuron, and interval of each assessed value. When assuming that the shape of the distribution of values was constant throughout the model (i.e. percentile was only included as a main effect), this third statistical model supports the original conclusion based on model 1: the effect of PTX on mEPSC amplitude was modulated by genotype (Fig. 4C, Genotype*Pharmacological treatment: F_2,75.72_ = 3.334, p = 0.0409). Interestingly, a more complex statistical assessment demonstrated that the interaction between genotype and pharmacological manipulation was statistically different at the five assessed percentiles of the measurement distribution (Percentile*Genotype*Pharmacological treatment: F_8,5064_ = 4.246, p < 0.0001). Post-hoc comparisons of this relationship revealed that the differences between genotype and pharmacological treatment could not be detected in the smallest amplitude mEPSCs (6^th^ percentile) but become progressively more pronounced with larger mEPSC amplitudes (26^th^, 50^th^, 74^th^ and 94^th^ percentiles). In conclusion, our novel approach provides a way to properly account for the hierarchical nature of repeated electrophysiological measurements that increases statistical power while providing the ability to statistically compare specific parts of distributions of measurement values between conditions.

## Discussion

The statistical analysis of most types of electrophysiological datasets is complicated by both the repeated nature as well as the hierarchical relationships between measurements. Traditional approaches to analyze these datasets typically rely on the implicit assumption that individual measurements are statistically independent [Aarts et al., 2014; Yu et al., 2022]. Our demonstration that the interevent interval and amplitude of EPSCs is clustered around the mean of individual neurons demonstrates that this assumption is violated and illustrates the importance of using statistical models that reflect the hierarchical nature of electrophysiological datasets. Beyond the similarity of measurements taken from individual neurons, our assessment demonstrates that the slice from which recordings were made influences measurements in some cases while other factors (time interval, mouse, and experimenter) barely influenced the outcomes. Based on these findings, we propose that analyses of most electrophysiological datasets should, as a minimum, incorporate the neuron and slice level into their statistical model.

Beyond assessments of average responses, comparisons of electrophysiological changes often involve a description of the distribution of values. Traditionally, these assessments rely on the two-sample Kolmogorov-Smirnov test. Unfortunately, the two-sample Kolmogorov-Smirnov test only compares a single value per distribution and does not take the hierarchy underlying measurements into account. Our novel approach identifying multiple values that together describe the distribution of values (i.e. the 6^th^, 26^th^, median, 74^th^, and 94^th^ percentile values) enables a more structured and reliable method to assess responses in specific parts of the distribution of outcomes. The selection of these specific percentiles is somewhat arbitrary, but we feel that a set of five values provides an optimum in the trade-off between describing the distribution more fully and being able to interpret potential differences in outcome. It is of course possible to select a different set of percentile values if an experiment requires a more (or less) precise description of the distribution of outcomes.

Despite the complexities presented by datasets of hierarchically organized and repeated electrophysiological measurements, these datasets also offer the potential of high statistical power when analyzed correctly. Our new EPHierStats approach is designed to use the power of parametric mixed-effects general linear models to enable the description of complex hierarchical relationships while analyzing repeated-measurements datasets at the level of individual measurements. By not reducing large sets of measurements to a single value per neuron and by incorporating biological replicates within each neuron and experimental condition, statistical power is maximized. A further benefit of the use of mixed-effects general linear models is that it allows assessments of more complex (interacting) relationships between groups and conditions. When analyzing such relationships, the most powerful analyses will be those that compare within-neuron manipulations since measurements from the same neuron cluster together, but a host of other between-neuron comparisons can also be assessed. Overall, the EPHierStats approach proposed here enables the description of hierarchical, repeated-measurements electrophysiological datasets in their full complexity and provides a powerful statistical tool to analyze the effects of experimental manipulations on the full distribution of measurement outcomes.

